# Selective maintenance mechanisms of seen and unseen sensory features in the human brain

**DOI:** 10.1101/040030

**Authors:** Jean-Rémi King, Niccolo Pescetelli, Stanislas Dehaene

## Abstract

Recent studies of “unconscious working memory” have challenged the notion that only visible stimuli can be actively maintained over time. In the present study, we investigated the neural dynamics of subliminal maintenance using multivariate pattern analyses of magnetoencephalography recordings (MEG). Subjects were presented with a masked Gabor patch whose angle had to be briefly memorized. We show with an unprecedented level of precision, that irrelevant sensory features of contrast, frequency and phase are only encoded transiently. Conversely, the relevant feature of angle is encoded and maintained in a distributed and dynamically changing manner throughout the brief retention period. Furthermore, although the visibility of the stimulus correlates with an amplification of late neural codes, we show that unseen stimuli can be partially maintained in the corresponding neural assemblies. Together, these results invalidate several predictions of current neuronal theories of visual awareness and suggest that visual perception relies on a long sequence of neural assemblies that repeatedly recode and maintain task-relevant features at multiple levels of processing, even under unconscious conditions.

## 1 Introduction

Conscious perception is often associated with the ability to hold a representation in mind. Indeed, the influence of invisible stimuli on behavior rapidly decreases with time. Furthermore, neuroimaging studies have repeatedly shown that invisible stimuli typically fail to evoke late and sustained neural responses, notably in fronto-parietal cortices [1]. Several models of visual awareness have consequently conjectured a strong link between the visibility of a stimulus and the maintenance of specific neuronal codes, through the coalition of coherent thalamo-cortical neuronal assemblies [2], recurrent processing within the cortex [3] or the sustained recruitment of the fronto-parietal networks [4].

The association between visual awareness and information maintenance has however been recently challenged. First, several groups have shown that an unseen stimulus can sometimes evoke late neuronal responses [5, 6, 7, 8, 10, 11]. Second, Soto and collaborators have recently shown in a series of behavioral experiments that a masked Gabor patch can be mentally maintained for several seconds, even when subjects report not seeing it [12, 13, 14]. Finally, functional Magnetic Resonance Imaging (fMRI) suggests that this unconscious visual maintenance depends on prefrontal activity [15, 13].

The current neuronal theories of visual awareness could offer ad-hoc explanations to account for the maintenance of these putatively unconscious stimuli. For example, the Global Neuronal Workspace Theory predicts that the maintenance of a neuronal module could remain unconscious if its output failed to be globally broadcasted across brain areas [4, 16]. Alternatively, the Recurrence Theory predicts that a long feedforward activation that fails to trigger recurrent processing could in principle provide a mechanism for an unconscious dynamic maintenance [3].

Testing these theoretical predictions requires identifying, the neural mechanisms responsible for the maintenance of unseen stimuli. Specifically, one needs to determine 1) whether the maintenance of unseen information is confined to early sensory regions or broadcasted to higher processing stages ([16, 4, 17]) and 2) whether the maintenance of an unseen stimulus depends on the sustained firing rate of a coding neuronal assembly (e.g. [18]), on the dynamic transmission of information across multiple modules (e.g. [19]), or both.

In the present study, we investigated with magneto-encephalography the neural mechanisms encoding and briefly maintaining low-level visual features, and tested how their recruitment varies as a function of the stimulus visibility.

## 2 Method

### 2.1 Stimuli & Protocol

Twenty young healthy adults were scanned with MEG (22±3 years old; 11 males, 18 right-handed). Subjects had normal or corrected-to-normal vision. Each experiment lasted for approximately one hour and was financially compensated. All subjects gave written informed consent to participate in this study, which was approved by the local Ethics Committee.

Each trial started with a brief and variably contrasted target Gabor patch (17 ms), subsequently masked by a radial sinusoid (117 ms, inter stimulus interval: 50 ms, Figure 1 a.). A probe Gabor patch was then presented for 67 ms, 800 ms after the onset of the target. The contrast of the target was pseudo-randomly varied between 0% (‘absent’ trials), 25%, 75% and 100%, whereas the contrast of the mask and of the probe was fixed to 100%. The orientation of the target pseudo-randomly varied between 15^°^, 45^°^, 75^°^, 105^°^, 135^°^, and 165^°^. The probe angle was tilted 30^°^ relative to the target angle; the direction of this tilt (clockwise or counter-clockwise) was varied pseudo-randomly.

**Figure 1:**
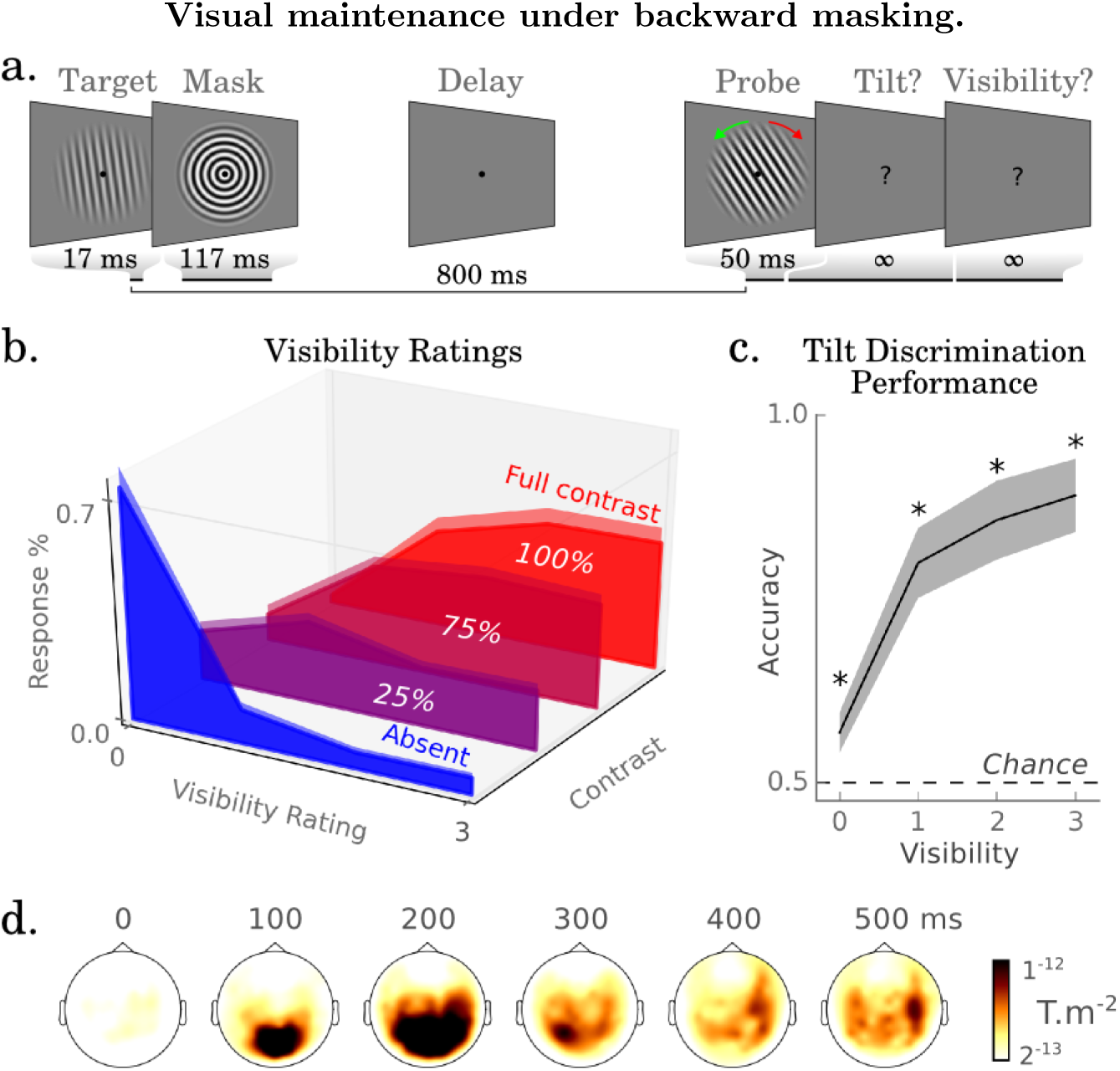
a. Subjects had to mentally maintain the orientation of a masked Gabor patch to compare it to a visual probe (clockwise or anti-clockwise tilt forced choice). At each trial, subjects reported the visibility of the target with a 4-point scale. b. The proportion of visibility reports for target absent trials and each level of contrast. c. Forced-choice tilt discrimination performance correlates with visibility reports, but nevertheless remains significantly above chance in trials reported as unseen. Error bars indicate the standard error of the mean (SEM). d. Norm of the planar gradiometers evoked by the target (present–absent trials).

Subjects made two successive decisions. First, they performed a forced-choice discrimination task, which consisted in indicating whether the probe was tilted clockwise or counter-clockwise to the target (index and middle finger of the left hand respectively). Subjects were then asked to report the visibility of the target (0: no experience of the target, up to 3: clear experience of the target, as defined by the ‘Perceptual Awareness Scale’ [20]), using the left index, middle, ring and little fingers of their right hand respectively. Subjects did 30 minutes of training before entering the MEG to ensure that they understood the task, and sensibly used all visibility ratings. Four subjects had to be excluded from the analysis because “unseen” or “clearly seen” reports were given less than 10 times across the experiment, or because the training phase had not been completed.

The phases of the target and of the probe randomly varied between -180° and 180°. The target spatial frequency pseudo-randomly varied between two possible values (30 and 35). The spatial frequency of the probe and of the mask was fixed to 32.5. The target, mask and probe stimuli had a fixed size of 16° of visual angle. Stimuli were presented on a gray background of a projector refreshed at 60 Hz, and placed 106 cm away from subjects’ head. Subjects were asked to keep their eyes opened and to avoid eye movements by fixating a dot continuously displayed at the center of the screen. Subjects performed a total of 840 trials, shuffled across five blocks of ∼12 minutes each. Pseudo randomization corresponds to a shuffled permutation of all conditions and was performed within each block.

### 2.2 Preprocessing

The preprocessing and statistical pipelines are available on github.com/kingjr/decoding_unconscious_maintenance, together with their modification history, several method tutorials, and an interactive preview of the results.

Magneto-encephalography recordings were acquired with an ElektaNeuromag^®^ MEG system (Helsinki, Finland), comprising 204 planar gradiometers and 102 magnetometers in a helmet-shaped array. Subjects’ head position relative to the MEG sensors was estimated with four head position coils placed on the nasion and pre-auricular points, digitized with a PolhemusIsotrak System^®^, and triangulated before each block of trials. Six electrodes recorded electro-cardiograms as well as the horizontal and vertical electro-oculograms. All signals were sampled at 1000 Hz, and band-pass filtered online between 0.1 and 330 Hz. Raw MEG signals were cleaned with the signal space separation [21] method provided by MaxFilter to i) suppress magnetic interferences ii) interpolate bad MEG sensors and iii) realign the MEG recordings into a subject-specific head position. The signals were then low-pass filtered at 30 Hz with a zero-phase forward and reverse Butterworth IIR filter (order=4), epoched between −300 ms and +1.200 ms relative to the target onset and baseline-corrected from −300 to −50 ms. Artefacted epochs were removed from the analysis after visual inspection. Epochs were finally down-sampled to 128 Hz.

The orientation of a Gabor patch ranges from 0 to 180°. To facilitate the circular analyses described below, we will refer to Gabor angle as the double of Gabor orientation. The phase of the Gabor patches was random. To facilitate the analyses and keep a consistent processing pipeline (i.e. the stratified k-folded cross-validation is only implementable with discrete values), continuous phases were digitized into 6 discrete evenly-separated bins.

Four large time regions of interest were used to simplify the results and maximize signal-to-noise ratio. The baseline, early, delay and probe time windows refer to time samples between −100–50 ms, 100–250 ms, 300–800 ms, and 900–1050 ms relative to the target onset respectively.

### 2.3 Statistics

Except if stated otherwise, statistical analyses were based on second order tests across subjects. Specifically, each analysis was first performed within each subject separately (across trials). We then tested the robustness of these effect sizes across subjects, using, whenever possible, non-parametric statistical tests, which tend to provide more robust, although potentially less sensitive statistical estimates. Except if stated otherwise, the reported effect sizes correspond to the mean effect size ± the standard error of the mean (SEM) across subjects; the p-values correspond to the second-order analyses obtained across subjects. Categorical and ordinal tests were based on two-tailed Wilcoxon and Spearman regression analyses respectively, as provided by the Scipy package [22]. Parametric circular-linear correlations were implemented from [23] and consisted in combining the linear correlation coefficients (R) obtained between the linear data (*x*) and the cosine and sine of the circular data (*α*): *R*_*sin*_ = *corr*(*x, sin*(*α*))

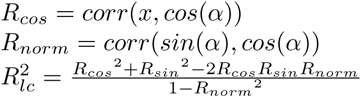

Where 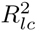 is the linear-circular correlation coefficient between x and *α*. Mass-statistical analyses, such as those used to test the significance of each channel at each time sample, or each estimator at each time sample, were based on cluster-based permutation analyses [24], using the default parameters of the MNE-Python *spatio temporal cluster 1samp test function*, which intrinsically corrects for multiple comparison issues. Uncorrected p-values are explicitly denoted as *p*_*uncorrected*_. Except if stated otherwise, analyses were only based on meaningful trials: for instance, the decoding of Gabor angle was solely based on target-present trials, and trials with missed decision responses were excluded from any analyses involving a decision factor.

### 2.4 Decoding

The multivariate estimators aimed at predicting a vector (*y*) of categorical (e.g. present versus absent), ordinal (e.g. visibility = 0, 1, 2, 3) or circular data (e.g. Gabor angle: 30^°^, 90^°^…, 330^°^) from a matrix of single trial MEG data (*X*, shape = *n*_*trials*_ × (*n*_*chans*_ × 1 *time sample*), Figure S8). Decoding analyses systematically consisted in i) fitting a linear estimator (*w*) to a training subset of X (*X*_*train*_), ii) predicting an estimate of *y* on a separate test set (*ŷ*_*test*_) and finally iii) assessing the decoding score of these predictions as compared to the ground truth (*score*(*y*, *ŷ*)).

#### Estimators

Each estimator made use of two processing steps. First, *X* was whitened by using a standard scaler that z-scored each channel at each time point across trials. Second, a linear support vector machine (SVM) algorithm was fitted to find the hyperplane (*w*) that maximally predicts *y* from *X* while minimizing a square hinge loss function. All SVM parameters were set to their default values as provided by the Scikit-Learn package (*C* = 1, *l*2 regularization, *tolerance* = 0.0001) at the exception of setting an automatic class-weight parameter that aimed at making the analysis more robust to potential class imbalance in the dataset. Three variants of estimators were implemented to deal with categorical, ordinal and circular data respectively. Categorical and ordinal tests were based on support vector classifiers (SVC) and support vector regressors (SVR) respectively. The SVC included a supplementary computational layer to generate probabilistic estimates instead of categorical predictions, by using a nested cross-validation based on Platt’s method [25]. Finally, a combination of SVR was used to perform circular correlations: two distinct SVR were fitted on *X* to respectively predict *sin(y)* and *cos(y)*. The predicted angle *(*ŷ*)* was estimated from the arctangent of the predicted sine and predicted cosine of the two SVR: *ŷ* = *artan2*(*ŷ*_*sin*_, *ŷ*_*cos*_).

#### Cross-validation

Each estimator was fitted on each subject separately, across all MEG sensors, and at a unique time sample (sampling frequency =128 Hz). In other words, for each analysis (decoding of Gabor angle, contrast, visibility report etc), we fitted *n*_time_ estimators on an X matrix (*n*_*trials*_ x *n*_*channels*_ x 1 time sample of MEG data) to robustly predict a vector *y* (*n*_*trials*_ x 1 categorical, ordinal or circular data). This analysis was performed with an eight folds stratified folding cross-validation, such that each estimator iteratively generated predictions on 1/8^th^ of the trials (testing set) after having been fitted to the remaining 7/8^th^ (training set) while maximizing the distribution homogeneity across training and testing sets (stratification).

#### Scores

Decoding analyses generated an *n*_*times*_ x 1 vector of probabilistic, ordinal or circular data *(ŷ)* that could be compared to the trials’ actual categorical, ordinal or circular value *(y)*. Categorical decoding was summarized with an empirical Area Under the Curve applied across all trials (AUC, range between 0 and 1, *chance* = 0.5). Ordinal decoding was summarized with a Spearman correlation R coefficient (range between −1 and 1, *chance* = 0). Circular decoding was summarized by computing the mean absolute difference between *ŷ* and *y* (range between 0 and *π*, *chance* = π/2). To facilitate visualizations, this ‘error’ metrics was transformed into an ‘accuracy’ metrics (range between −π/2 and π/2, *chance* = 0).

#### Time regions of interest

Except if stated otherwise, the decoding scores obtained for a large time window of interest was generated by i) averaging the decoding predictions across the selected time samples at the single trial level, ii) computing the unique resulting score for each subject and iii) performing a univariate categorical or ordinal test across subjects. Averaging of circular data (e.g. decoded angle of a Gabor patch) was performed in the complex space: 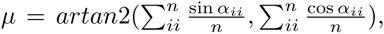 where *α*_ii_ is the angle at trial *ii, n* is the number of trials, and *μ* the average angle.

#### Temporal generalization

Time-resolved decoding analyses are a specific case of temporal generalization analyses where the estimators are fitted, tested and scored with a unique time sample (Figure S9). Each estimator fitted across trials at time *t* can also be tested on its ability to accurately predict a given trial at time *t’*, so as to estimate whether the coding pattern is similar between *t* and *t’*. When applied systematically across all pairs of time-samples, this analysis results in a square generalization-across-time (GAT) matrix, where the y-axis corresponds to the time at which the estimator was fitted, and the x-axis to the time at which the estimator was evaluated.

All decoding analyses were performed with the MNE-Python [26] and Scikit-Learn packages [27]. The first-order decoding analyses have been integrated to the MNE-Python package under the *TimeDecoding and GeneralizationAcrossTime* classes.

### 2.5 Topology analyses

Two-dimensional graphs were generated from temporal generalization analyses. The nodes and edges of the graph correspond to the training and testing times of the GAT matrix respectively (Figure 6). Indeed, each training time corresponds to a specific estimator, itself isolating a linear combination of MEG sensors, and thus a specific, possibly distributed, neural assembly. The similarity of these neural assemblies can thus be estimated from the co-activation of their respective estimators [28].

Each graph was based on the average of decoding scores obtained across subjects. This connectivity matrix was normalized by the maximum decoding score thresholded according to the second order statistical significance obtained with cluster-permutation analyses, and rectified (negative weights correspond to topographical inversions and thus imply the same neural regions than positive weights). The sizes of the nodes are proportional to the decoding scores obtained at the time at which each estimator was trained. The two-dimensional position of the nodes was initialized around a circle, and iteratively re-estimated with the Fruchterman-Reingold force-directed algorithm until reaching a local minimum (*n*_*iterations*_=100) provided in the NetworkX package [29].

The resulting 2D plot attempts to summarize a complex hyper-dimensional graph, and is thus necessarily simplistic and partially arbitrary. Different approaches, such as multidimensional scaling and spectral embedding would lead to slightly different 2D layouts. The critical features of the graph therefore relate more to the topology of the network than the exact distance that separate the nodes.

## 3 Results

### 3.1 Behavioral evidence of a weak maintenance of unseen stimuli

We first quantified the extent to which subjects were able to detect the masked Gabor patches (target), maintain its orientation and compare it to a subsequent probe (Figure 1 a). Subjects’ visibility ratings varied across the four-point visibility scale (0: unseen, 3: clearly seen). Absent trials were generally reported as unseen (visibility=0/3: 74±6%) and present trials were generally reported with one of the three other visibility ratings (visibility > 0/3: 93±2%), leading to a detection d’ of 2.73±0.32 (Figure 1 b). This result confirms that subjects meaningfully used subjective visibility reports. Forced-choice discrimination performance (the ability to determine whether the probe was oriented clockwise or counter-clockwise to the target) was relatively high (85±5%, chance=50%), and strongly varied as a function of visibility (R=0.79±0.10, p<0.001), indicating that subjects adequately estimated their ability to detect the target. To our surprise, discrimination performance did not appear to systematically increase with the contrast of the target (R=0.17±0.14, p=0.194), although this effect may be under-powered by the fact that the contrasts only varied between three possible values. Importantly, target reported as unseen were discriminated slightly above chance level (accuracy: 58±5%, p=0.036; d’=0.20±0.09, p=0.006, Figure 1 c). These behavioral results suggest that subjects were weakly but significantly able to maintain and compare the orientation of a target stimulus to that of a probe presented 800 ms later, even when they reported not seeing the target.

### 3.2 The brain automatically encodes all sensory features in parallel

A series of regression analyses was applied for each sensor (univariate topography analyses) or across all sensors at once (decoding analyses) to track, at each time point, the neural responses specifically coding for each sensory and decision feature.

Comparing the event related fields (ERF) evoked in trials with a target and a mask, to trials with a mask but no target (’absent trials’) confirmed that the visual target elicited a strong focal response on centro-posterior MEG channels from between ∼80 and 250 ms after the onset of the target (average decoding scores 100–250 ms: AUC=0.91±0.01, p<0.001, Figure 1 d).

These early ERFs specifically encoded target orientations. Indeed, linear circular correlations between the ERF and the target angles revealed a focal response over posterior channels from ∼90 ms and the corresponding decoding scores were relatively low but strongly significant during this early time window (100–250 ms: 0.060±0.007 rad., p<0.001, Figure 2). Similar analyses applied onto probe orientations revealed analogous ERF and decoding results at the notable exception of a much higher signal-to-noise ratio (900–1050 ms after target onset, i.e.100–250 ms after probe onset: 0.111±0.008 rad., p<0.001, Figure 2).

**Figure 2:**
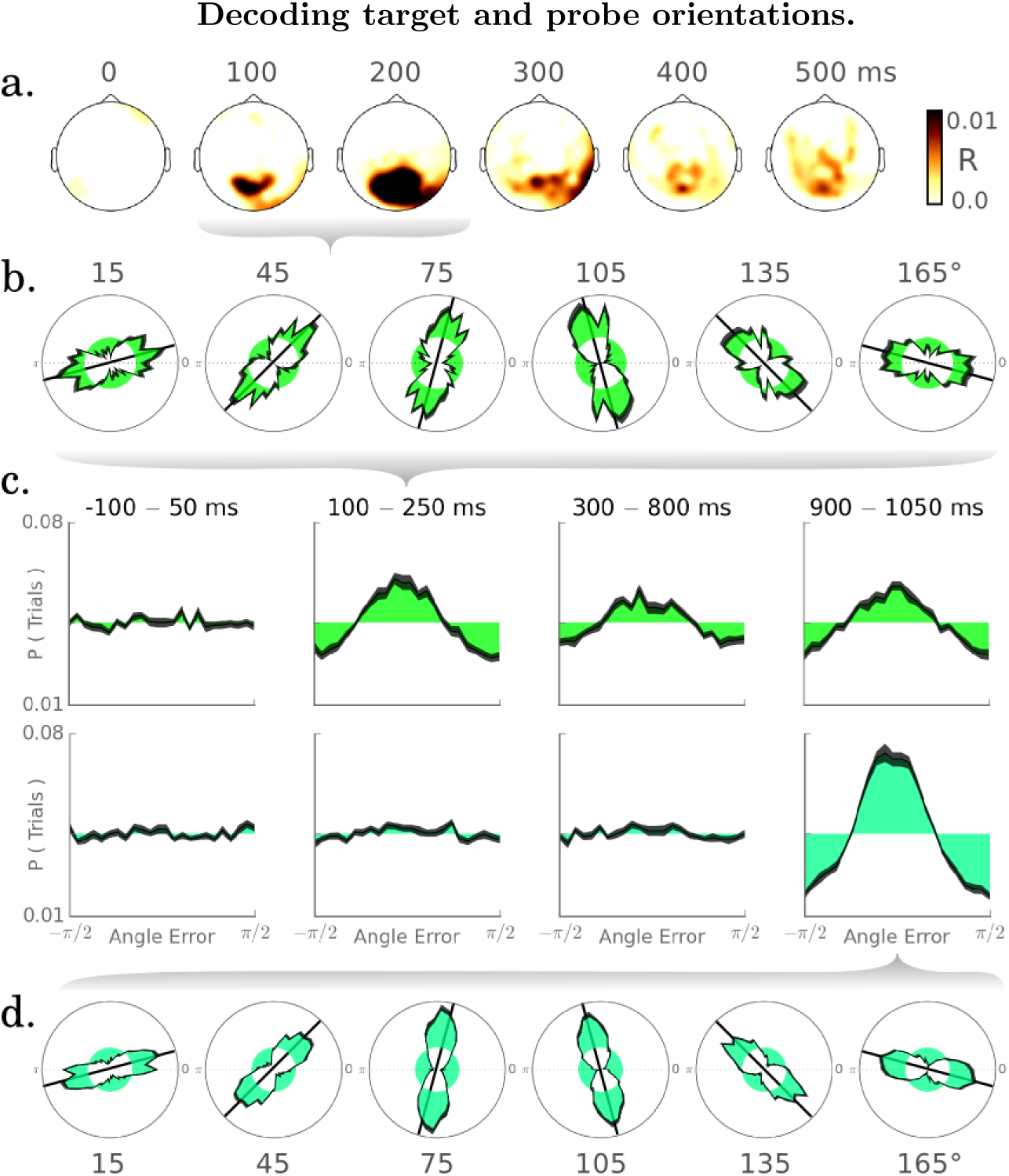
a. The linear-circular correlations between MEG signals and target angle peak around ∼100-250 ms over centro-posteriors regions. Topographies depict the combined effect sizes for each pair of gradiometer. b-d. Decoded target (green) and probe angles (turquoise) for each possible stimulus. c. Histograms of angle errors obtained in four time regions of interest locked to target onset. Error bars indicate the SEM across subjects and filled areas indicate chance level.

The contrast, spatial frequency and phase of the target (Figure 3) also appeared encoded in these early brain responses: univariate analyses consistently revealed correlations between the early posterior responses and each of these sensory features. Although some of these effects vanished after correction for multiple comparisons, decoding analyses confirmed that the contrast (R=0.17±0.01, p<0.001) and the spatial frequency of the target (AUC=0.53±0.01, p=0.005), as well as the phase of the probe (0.054±0.012 rad., p<0.001) could be decoded significantly above chance between approximately 100 and 250 ms after the onset of the corresponding stimulus. The decoding of the target phase did not reach statistical significance, but additional analyses showed that the estimators fitted to the probe phase significantly predicted the target phase between 148 and 202 ms after target onset (0.024±0.006 rad., p<0.001). The phase of the target thus appears to be decodable from the MEG response, but the signal is so weak that the default parameters used to fit the initial estimators were visibly suboptimal to robustly capture this effect.

**Figure 3:**
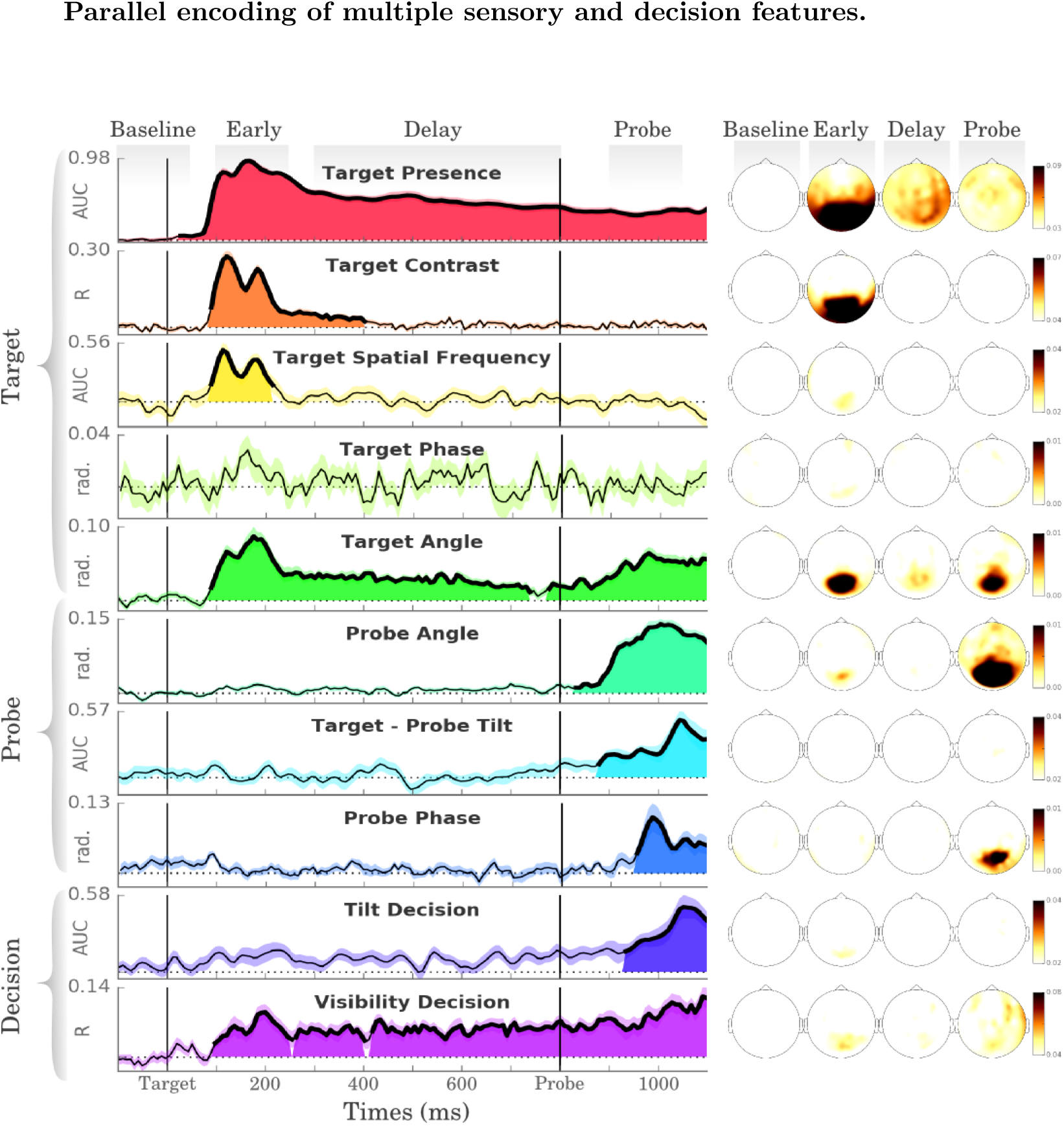
Time course of decoding performance for each sensory and decisional feature. Filled areas and thick lines indicate significant decoding scores (cluster corrected, p < 0.05) and dotted lines indicate theoretical chance level. For each feature, the corresponding coding topographies are depicted with combined gradiometers in each of the four time windows of interest.

Overall, these results demonstrate that all of the sensory features presently manipulated are simultaneously and automatically encoded in early brain responses.

### 3.3 Task-relevant visual features are selectively maintained

While the task-irrelevant sensory features of contrast, spatial frequency and of phase were not detectable beyond 250 ms in the MEG activity, the task-relevance features of presence and orientation as well as the visibility of the target remained decodable during the entire epoch (Figure 3).

Specifically, the decoding scores of the target presence were significantly above chance during the delay period (300–800 ms: AUC=0.73±0.02, p<0.001) as well as after probe onset (900–1050 ms: AUC=0.66±0.02, p<0.001, Figure 3, red), and was characterized by spatially distributed MEG response from ∼250 ms onward. Similarly, the decoding of target orientation was low but remained significant from ∼250 ms (300–800 ms: 0.025±0.003 rad., p=0.005). The corresponding univariate ERFs only reached statistical significance after the probe onset, suggesting that the anatomical substrates recruited during the delay time period may have been too variable across subjects to be detectable with conventional group analyses across sensors. Additional control analyses confirmed that the decoding of the target after probe onset could not be solely explained by the correlation between the angles of these two stimuli (Supplementary Figure S11).

Interestingly, we observed that visibility, a subjectively-defined but task-relevant visual feature was also detectable throughout the retention period. Indeed, the decoding of visibility decisions was low but sustained from ∼100 ms up to the end of the epoch (100–250 ms: R=0.06± 0.01, p=0.007, 300–800 ms: R=0.05±0.01, p=0.002, 900–1050 ms: R=0.08±0.01, p=0.002). By contrast, the decoding of forced-choice discriminations was only significantly above chance around probe onset (900–1050 ms: AUC=0.54±0.01, p=0.002, Figure 3).

Overall, these results suggest that, the brain automatically encodes all visual features in parallel around 100-250 ms, but then only processes and maintains those relevant to the task (Figure S10).

### 3.4 The maintenance of unseen sensory information can be tracked over time

To investigate whether the maintenance of unseen visual information could be detectable in the MEG results, we separately analyzed seen and unseen trials, as defined in a conservative manner (seen=rating 3/3, unseen= rating 0/3).

Decoding the presence of the target was significant in both seen and unseen conditions during the early time window (AUC: seen 0.91±0.01, unseen: 0.87±0.02, both p<0.001; for clarity purposes, we report the AUC of present trials as compared to all absent trials) as well as during the retention period (AUC: seen: 0.74±0.02, unseen: 0.65±0.02, both p<0.001, Figure 4. Top left).

**Figure 4:**
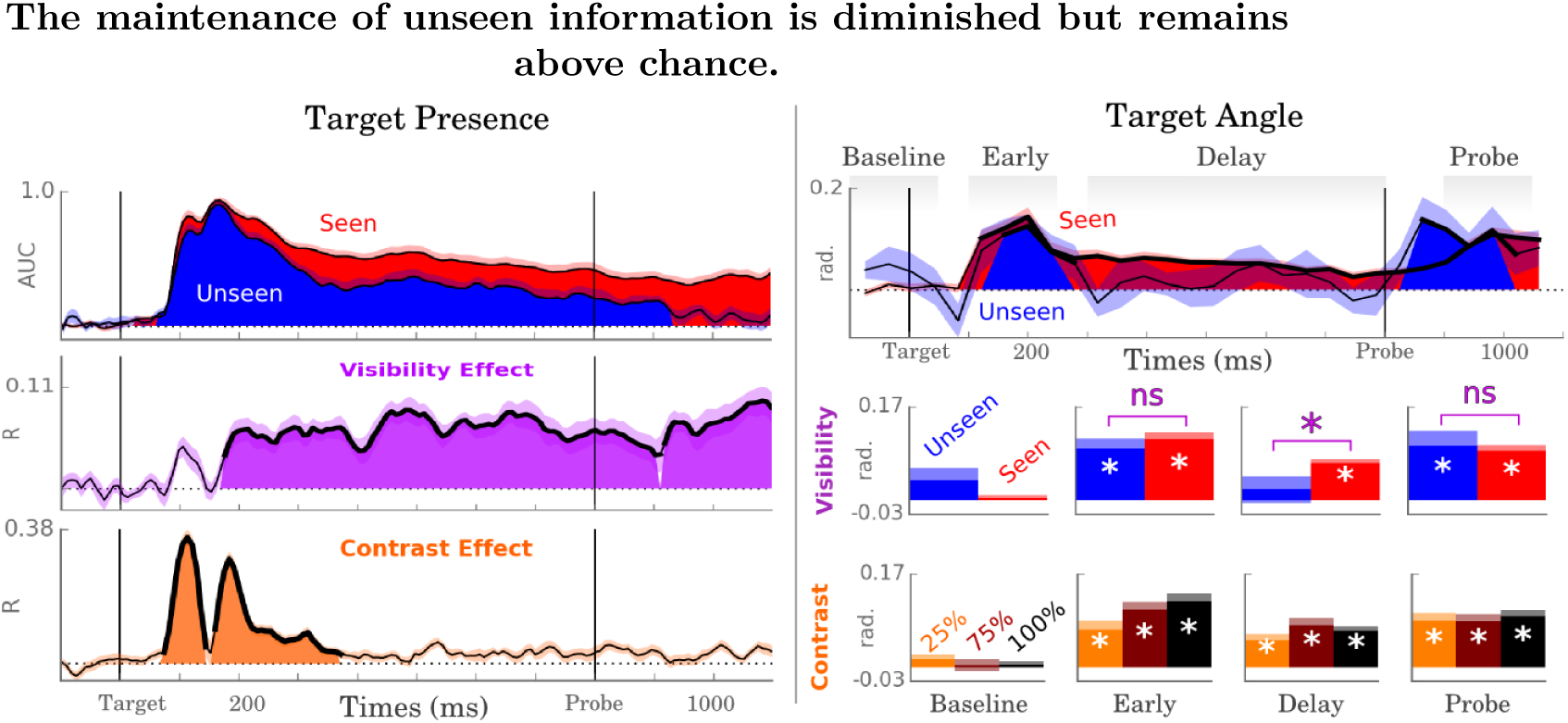
Left. Decoding subscores of the target presence as a function of time and visibility (red: seen, blue: unseen; for clarity purposes, the decoding scores are estimated against all absent trials). The decoding scores of target presence were specifically modulated by the contrast of the target contrasts from ∼80 to 300 ms (orange) and by visibility ratings from ∼180 ms (purple). Right. Decoding subscores of the target angle as a function of time and visibility. The time courses are down-sampled for visualization purposes. Target angle decoding scores were modulated by target contrasts during the early and the probe time periods, and by visibility ratings during the delay period.

The decoding scores of seen and unseen orientations were noisier than those of presence decoding, but presented a similar overall picture to the one observed above with the presence decoding (Figure 4 Top right). Specifically, the orientations of both unseen and seen trials could be decoded shortly after the target onset (seen: 0.111±0.13 rad., p<0.001, unseen: 0.085±0.018 rad., p<0.001). These seen and unseen decoding scores did not appear significantly different from one another during these two time periods (100–250 ms: p=0.117; 900–1050 ms: p=0.576). Contrarily a significant difference was observed between seen and unseen trials during the delay period (seen-unseen: 0.047±0.025 rad., p=0.033). Unlike seen trials (0.067±0.008 rad., p<0.001) the decoding of unseen orientations did not reach statistical significance during the delay time period (0.019±0.024 rad., p=0.332), but only reached significance after probe onset (0.197±0.055, p=0.006). Additional analyses confirmed that these late codes could not be solely explained by the correlation between the target and the probe orientations (Figure S11).

Overall, these results confirm that visual information can be partially maintained even when the stimulus is reported as unseen.

### 3.5 The maintenance of sensory features is specifically affected by visibility decisions

In spite of being statistically detectable, the maintenance of unseen stimuli was largely deteriorated as compared to that of seen stimuli. Indeed, the decoding of the target presence positively correlated with subjective visibility ratings (0-3) from ∼180 ms (Figure 4 b., R=0.066±0.008, p<0.001), and seen orientations were better decoded than unseen ones (seen-unseen: 0.047±0.025 rad., p=0.033). This modulation of presence and orientation decoding scores as a function of visibility was specific to the delay period, and was not observed before ∼200 ms.

The modulation of sustained brain activity was specific to subjective criteria, and was remarkably independent from the objective stimulus contrast. Indeed, the contrast of the target modulated the early decoding scores of presence (R=0.187±0.014, p<0.001) and orientation (R=0.55±0.13, p=0.002), but rapidly stopped influencing both of these codes during the delay period (Figure 4 c). The modulations of decoding score by subjective visibility and target contrast factors were significantly different from one another during the early (Δ*R* = 0.149 ± 0.014, *p* < 0.001) and delay time windows (Δ*R* = −0.035 ± 0.009,*p* = 0.003), confirming the temporal specificity of these two factors. Note that we did observe a weak correlation between orientation decodability and target contrast after probe onset (R=0.325±0.129, p=0.0369) but the low confidence of this unexpected effect, together with the absence of such trend in the stronger presence decoding results, suggest a family-wise error.

Overall, these results suggest that the early encoding of the target features was performed independently of visibility, and was solely modulated by stimulus contrast. Conversely, the late processing stages were strongly modulated by subjective visibility (confirming previous reports [6, 30, 10]) but not by the objective stimulus contrast.

### 3.6 The maintenance of seen and unseen information depends on a long sequence of distributed neural assemblies

The temporal profile of decoding scores quantifies the amount of decodable information irrespective of the dynamics of its underlying neuronal substrate. To investigate the functional organization underlying the encoding and the maintenance of the target representation, we therefore applied temporal generalization analyses by testing whether each estimator trained at a given time point could generalize to all other time points. This analysis results in a generalization-across-time (GAT) scoring matrix, where the *y* and x-axes correspond to training and testing times respectively, and where the diagonal directly corresponds to the decoding scores presented in the previous sections (Figure S8). Different functional organizations result in specific GAT matrices and can thus characterize the functional organization underlying information maintenance (Figure S9). For example, a sustained neural activity typically results in a square component, whereas sequential neural activations are typically characterized by a diagonal component.

The empirical GAT matrices of each decoded features were typically characterized by i) an early reversal of the coding activity around ∼200 ms and ii) a relatively ‘thin’ early ‘diagonal’ component and iii) a relatively ‘thick’ late ‘diagonal’ component (for the relevant features only). Such diagonal component indicates that the ability of each estimator to generalize over time is relative brief as compared to the time period during which the corresponding features can be decoded. Diagonal components therefore indicate that the neuronal assemblies, isolated by each estimator, code their respective features only in a transient manner. Several weak off-diagonal components were also observed: orientation estimators trained around 200 ms weakly anti-generalized from ∼500 to ∼750 ms (0.01 < p < 0.05) and presence estimators trained around 250 ms generalized up to 600 ms, whereas later estimators weakly anti-generalized from 700 ms (0.01 < p < 0.05). These off-diagonal components suggest that the corresponding neural assemblies were sustained or reactivated. However, these effects were statistically weak and variable across presence and orientations estimators, and therefore remain difficult to interpret.

The functional organization underlying these temporal generalization analyses can be analyzed as a graph (Figure 6 a-c), where each node corresponds to an estimator and each edge is defined as the ability of the corresponding estimator to generalize to other time points. The overall layout depends on the number and on the weight of their edges, and should thus be interpreted in terms of overall topology, and not strictly on spatial location (see method). Overall, the graph visualizations illustrate i) the strong diagonal component of GAT matrices as a long chain of nodes and ii) the off-diagonal components as edges that curve of this chains.

**Figure 5:**
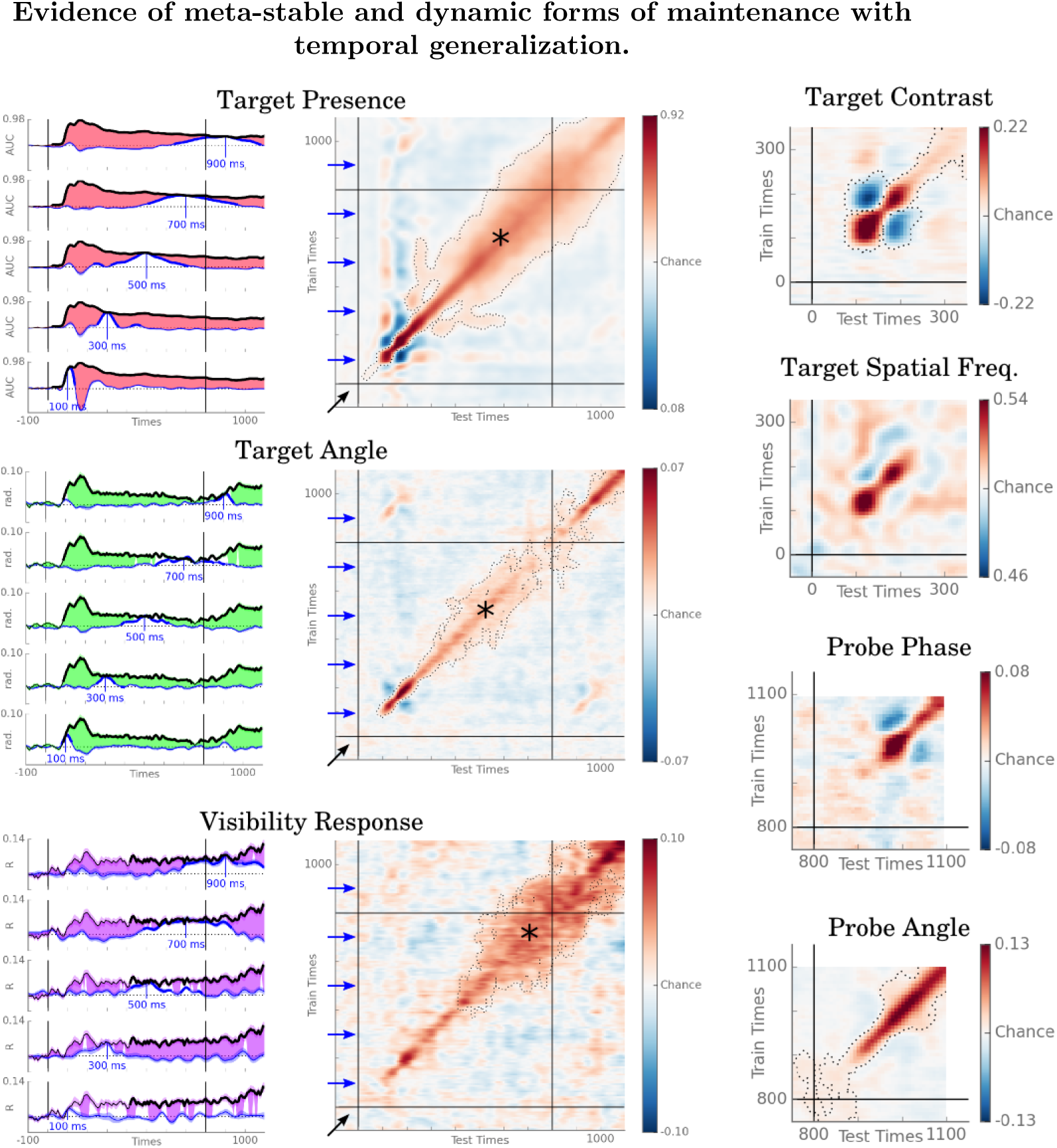
Temporal generalization matrices computed for the most relevant sensory and decisional features. Significant clusters (p < 0.01) are contoured with a dashed line. Diagonal clusters indicate that the coding MEG response changes over times. Below chance generalizations (blue) indicate a reversal of a brain activity (e.g. P1 / N1 couple). The time of course of decoding performance of five estimators (blue lines) trained at 100, 300, 500, 700 and 900 ms and compared against the diagonal of the GAT matrix (black lines). Filled areas indicate significant difference between the diagonal and the selected estimator; thick lines indicate scores significantly different from chance level.

**Figure 6:**
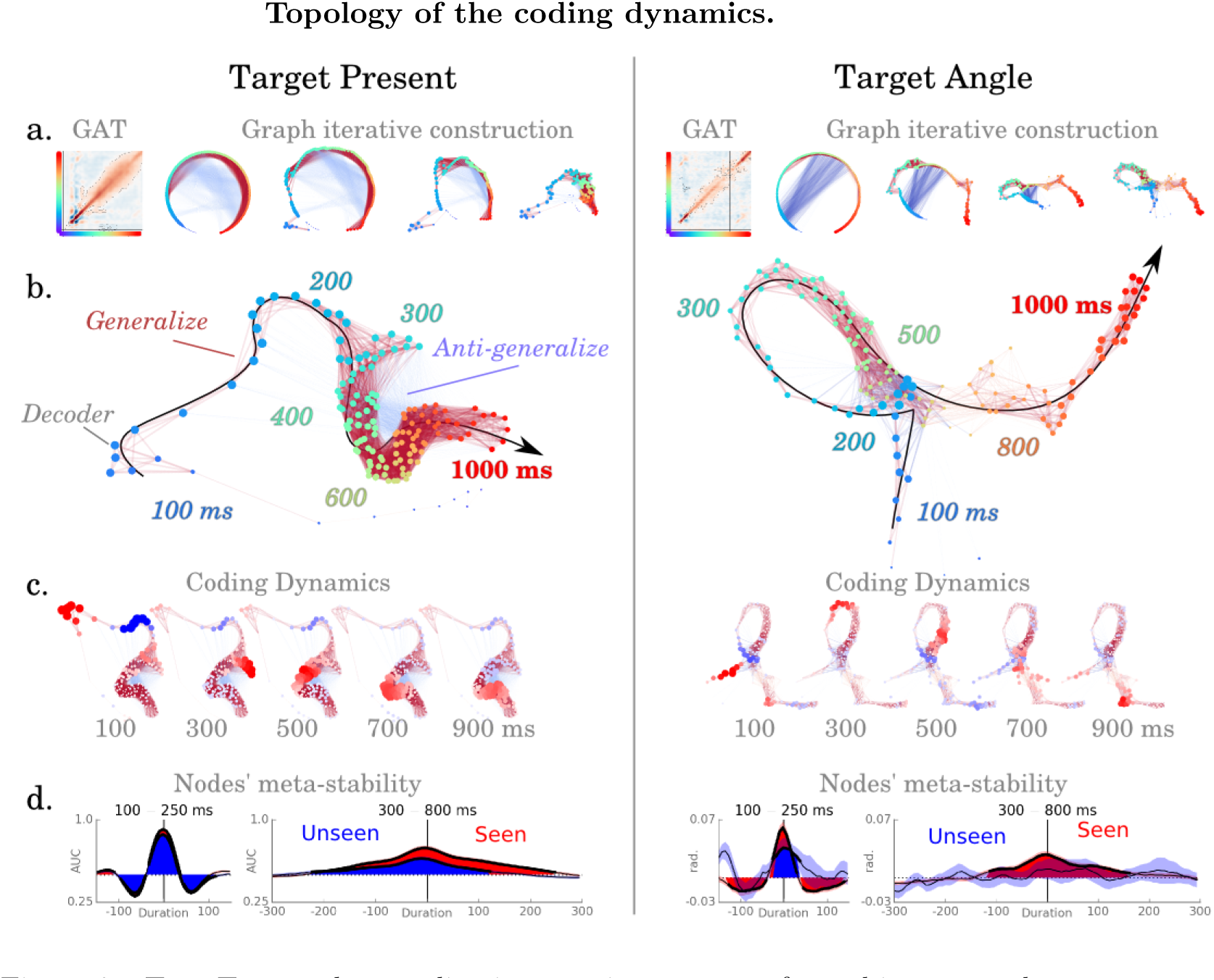
Top. Temporal generalization matrices are transformed into a graph of in which each node represent an estimator trained at a particular time point (color coded with a rainbow spectrum: blue=0, red=1000 ms) and each edge represent the extent to which each decoder generalize across time point (red=above chance generalization, blue=anti-generalization). The topology of the graph is summarized in 2D by attracting co-activated estimators towards one another. Middle. The dominating diagonal component of the GAT matrices translates in a long chain of nodes for the decoding of target presence (left) and target orientation (right). Off-diagonal generalizations, indicating the reactivation of early processing units, tend to curve this long chain. Coding dynamics show the decoding score of each estimator at five time samples. Bottom. The mean temporal generalization of early (100-250 ms) and late estimator (300-800 ms) reveals the meta-stability of each processing stage. Early processing stages are transient and independent of visibility for both presence and orientation codes. Late processing stage are maintained for several hundreds of ms in seen trials (red), but are reduced (target presence: ∼100 ms) or non-significant (target orientation) in unseen trials (blue).

### 3.7 The meta-stability of neural assemblies correlates with visibility, but remains detectable in unseen trials

Temporal generalization analyses revealed a dominating diagonal component, which suggests that the maintenance of sensory information is mainly performed by a long sequence of neural assemblies. To test whether subjective visibility correlates with the stability of particular neural activities [18], we quantified the ability of presence and orientation estimator to generalize over time as a function of visibility reports (Figure, 6 d.). Early estimators were typically more transient (presence: 42±1 ms, orientation: 96±19 ms) than late estimators (presence: 292±24 ms, p<0.001, orientation: 159±19 ms, p=0.010). Furthermore, the duration of early estimators did not significantly differ between seen (presence: 39±1 ms; orientation: 30±28 ms) and unseen trials (presence: 39±1 ms, p=0.072; orientation: 43±47 ms, p=0.446). By contrast, the duration of late estimators was significantly shorter for unseen (presence: 225±27 ms; orientation: 31±35 ms) than for seen trials (presence: 287±25 ms, p=0.006; orientation: 194±32 ms, p=0.0585), and the interaction between these early and late durations was significant (presence: p < 0.001; orientation: p=0.0239). Overall, these results confirm that i) early and late signals are typically transient and sustained respectively ii) that visibility partially and specifically correlates with the maintenance of late neural signals but iii) these results importantly show that the neural assemblies recruited after ∼250 ms can maintain their activity in unseen trials.

## 4 Discussion

### 4.1 Unseen but task-relevant sensory features can evoke late and sustained representations

Our behavioral results confirm previous findings showing that an unseen stimulus can be maintained over time [13], and further demonstrate that unseen but relevant sensory features can be decoded and tracked from subjects’ MEG activity during a brief retention period.

Although unseen representations appear significantly deteriorated as compared to seen ones, several results suggest that these sustained codes depend on an active maintenance mechanism rather than on the passive residues of sensory responses. Indeed, while all sensory features were decodable early on after the onset of the stimulus (< ∼250 ms). Only the features relevant to the task (presence, orientation and visibility of the stimulus) remained significantly maintained during the retention period (>∼300 ms). These results therefore suggest a dissociation between the automatic encoding and the selective maintenance of visual features. Topographical analyses strengthen this dissociation by consistently showing a focal response over visual regions and a spatially distributed set of responses for the early and the delay time windows respectively.

This two-stage processing hypothesis fits with recent studies decoding visible Gabor patches. Indeed, whereas Wolff et al. recently showed that a to-be-maintained Gabor patch [31] can be decoded from EEG for approximately one second, Ramkumar et al. [32], who decoded the orientation of task-irrelevant visible Gabor patches with MEG, only observed significant decoding scores for < 300 ms, despite higher signal to noise ratio than the one obtained in the present study.

Together, these results therefore suggest that an active and selective neuronal mechanism is recruited from ∼200 ms to selectively process and maintain the visual feature of orientation which is relevant to the task, even when the corresponding stimulus remains unseen.

### 4.2 Residue of unseen stimuli can be selectively broadcasted

To test current neuronal models of visual awareness, we assessed with temporal generalization analyses whether unseen stimuli could be maintained locally and/or dynamically, and whether they could be broadcasted in a similar way to seen stimuli. The resulting matrices were mainly dominated by a diagonal pattern lasting for more than 800 ms. Such long diagonal patterns, typical of a sequential processing ([28], Figure S9), suggest that the corresponding neural representation is coded i) transiently (each neural assembly is active for a shorter time window than the length of the diagonal), ii) redundantly (the same family of linear regressors decode the activity from different (sets of) brain regions) and iii) sequentially (the estimators are activated one after the other).

Similar long diagonal patterns have been repeatedly observed with EEG or MEG in recent visual studies [10, 31, 33, 34, 36, 37] as well as auditory and olfactory studies [38, 39]. Together, these diagonal patterns strengthen a series of anatomical and functional studies unraveling the hierarchical organization of the cortex [40, 41, 42]. Our results supplement this view by suggesting that visual information can be broadcasted across the cortical hierarchy even when the stimulus remains subjectively invisible. Indeed, unseen stimuli were neither characterized by an early disruption of the diagonal pattern as expected from a lack of broadcast, nor by a sustained activation of early processing stages, as expected from a maintenance mechanisms confined to early visual cortices. On the contrary, our results suggest that the dynamical maintenance and the broadcast of visual information were qualitatively similar across visibility conditions (Figure 7).

**Figure 7:**
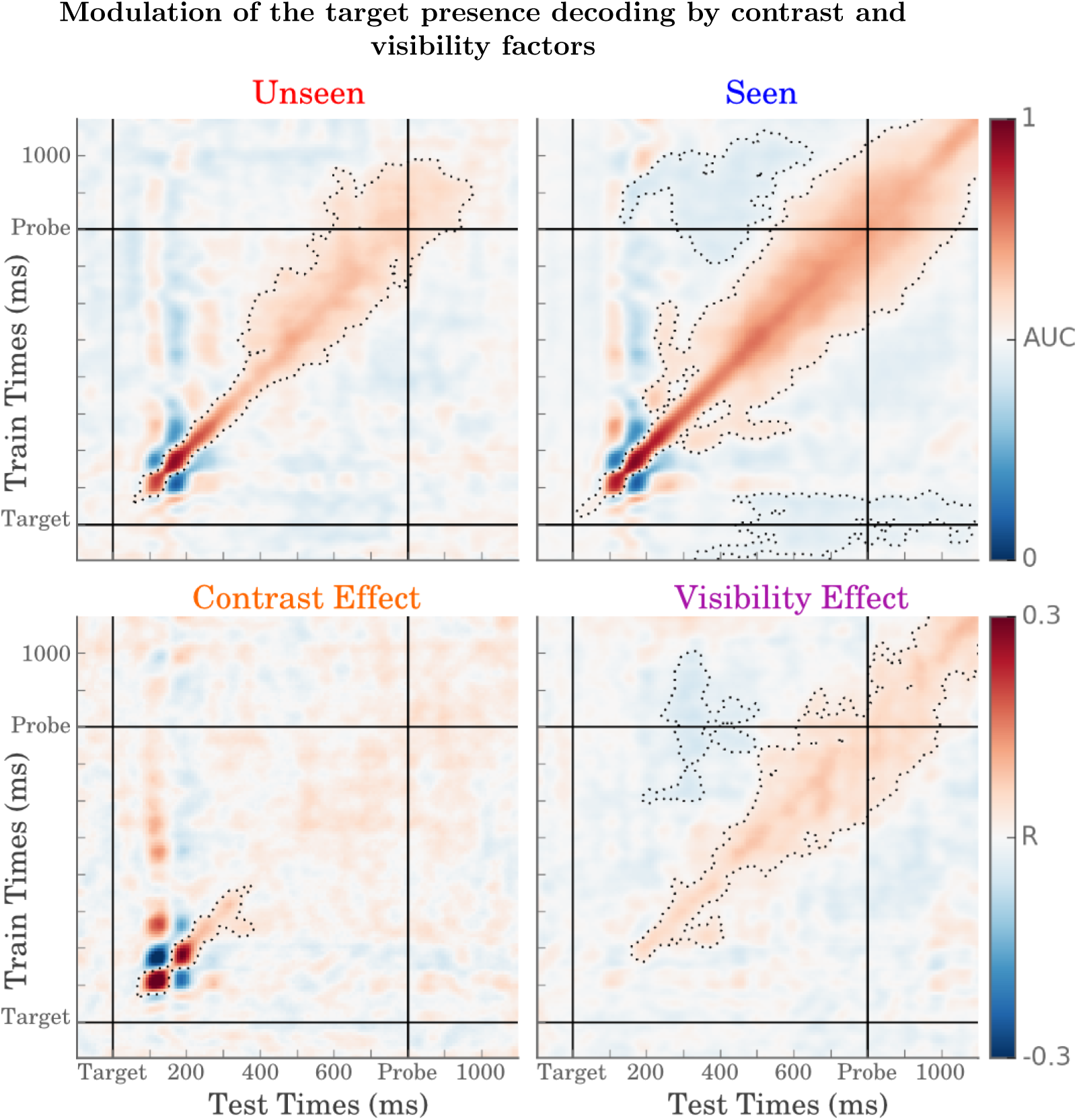
Top. Analyzing the temporal generalization matrices of target presence in unseen and seen trials separately confirmed that unseen information i) is encoded in all processing stages preceding probe onset, and ii) leads to partially meta-stable activity in late processing stages (i.e. thickening of late components of the diagonal). Bottom. Decoding scores of target presence were specifically modulated by the contrast (left) and the visibility (right) of the target during early (< 300ms) and late time windows (>200 ms) respectively. Significant GAT clusters (p <.01) are contoured.

### 4.3 Unconscious representations can be weakly meta-stable

Could this long diagonal reflect a purely feedforward phenomenon and thus support Recurrent Theory [3]? Our results demonstrate that the width of the temporal generalization diagonal increases from a few dozens of milliseconds before ∼250 ms to a few hundreds of milliseconds thereafter. As this width depends on the duration of activation of the corresponding neural assemblies, our results suggest that visual stimuli first recruits a series transient processing stage and subsequently evoke meta-stable responses from ∼250 ms. This observation fits with the ‘ignition’ phenomenon observed with scalp (e.g. [6, 30, 10] and intracranial recordings [43, 44, 3]).

Critically however, the width of the diagonal observed in the unseen conditions also thickens after ∼250 ms, suggesting that meta-stability is not unique to subjectively visible stimuli. Although this result challenges Recurrence Theory, it remains compatible with its original empirical support. Indeed, the intracranial studies distinguishing feedforward and recurrent processing as a function of visibility strictly focused on early visual regions [3, 4]. On the contrary the present MEG study investigates meta-stable neural activity across a wide variety of brain regions, and in fact suggests that early visual responses detectable with MEG are brief in both seen and unseen conditions. In the future, intracranial recordings of associative cortices could help overcome the spatial resolution of MEG and directly investigate the existence of recurrent processing during unconscious working memory task. Such approach could also help distinguishing silent working memory mechanisms from neuronal activity invisible to MEG recordings because of unaligned axons, or components not tangential to the MEG sensors [45].

### 4.4 Conscious perception is a distributed decision independent from maintenance mechanisms

Overall, the present study demonstrates that a low-level stimulus reported as completely unseen can be partially broadcasted to multiple brain areas, and be locally maintained in late processing stages. The present findings call for a partial revision of the neuronal mechanisms of visual awareness, but nevertheless remain profoundly compatible with their original empirical support. In particular, the time courses of the MEG topographies courses largely confirm that i) unseen visual features first evoke an early and automatic neural response that depends on the objective but not on the subjective properties of the stimulus, whereas ii) later neural responses are specifically modulated by subjective visibility, which consequently makes the detection of late unconscious processing difficult to detect.

Such unconscious broadcast and maintenance mechanisms could directly account for the existence of late and sustained unconscious brain responses [5, 6, 7, 8, 9, 10, 11, 12, 13, 14], and yet continue to support the idea that subjective visibility is a perceptual decision computed after ∼200 ms by a large network of neural modules [46, 17, 47, 48, 4, 49]. In this view, many neural modules can maintain residual sensory evidence long after an invisible stimulus is gone, but this lingering information, in a given trial, remains too similar to noise for subjective inference processes to conclude to the presence of the stimulus.

## 6 Supplementary Figures

**Figure S8:**
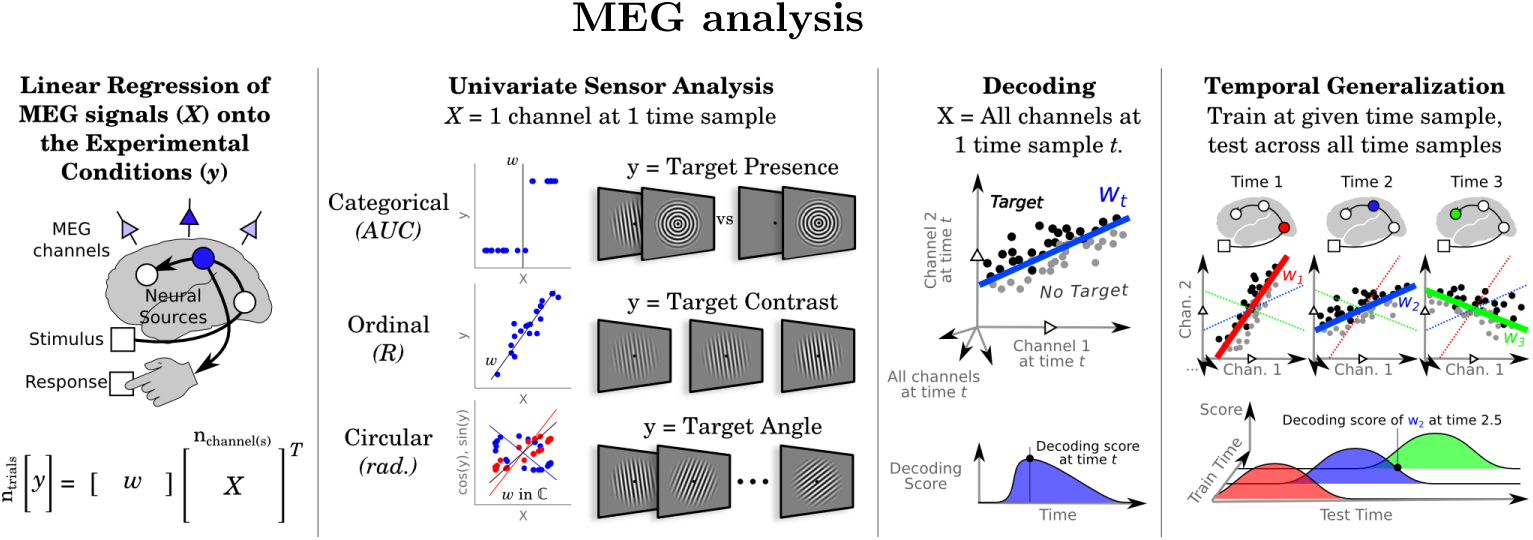
All MEG analyses aimed at identifying the regression coefficient(s) (*w*) of the single or multi-channel MEG signals (*X*) to an experimental condition (*y*) for each subject and at each time point separately. The vector *y* represents one sensory or decision features at each trial. Our ability to regress the MEG signals to categorical, ordinal and circular variables is summarized with an Area Under the Curve (AUC), R coefficient and angular error respectively. Note that for circular data, the linear function consisted in fitting the sine and cosine of y. In multivariate analyses, each trial *xin*X can be interpreted as a point in space where each dimension corresponds to an MEG sensor recorded at a given time. Decoding scores across time are estimated by repeatedly applying such multivariate analysis at different time points. Finally, temporal generalization analyses consist in quantifying the extent to which each estimator, fitted at a given time point, is able to generalize to other time points and typically result in a 2D generalization-across-time (GAT) matrix in which the y-axis corresponds to the time at which the estimator was fitted, and the x-axis, to the time at which the estimator was scored across trials.

**Figure S9:**
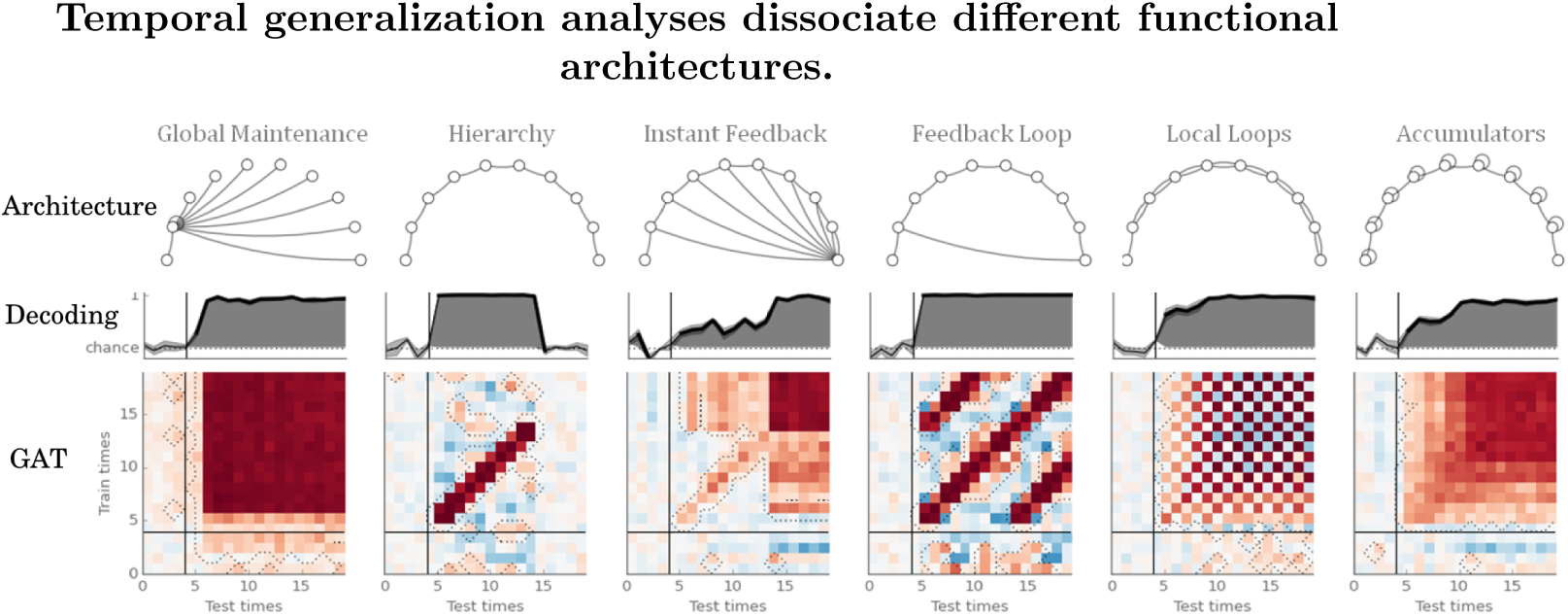
Different neural architectures can maintain information over time. For example, a unique early processing stage can directly maintain its activity while broadcasting its information to other areas (’Global maintenance’). Alternatively, the neural activity can be sequentially recoded by a long hierarchical network (’Hierarchy’). Other architectures based on sustained or dynamical feedback and/or horizontal connections can form different families of neural networks. While all of these networks can present similar, and potentially identical decoding scores across time, their GAT patterns are often distinguishable.

**Figure S10:**
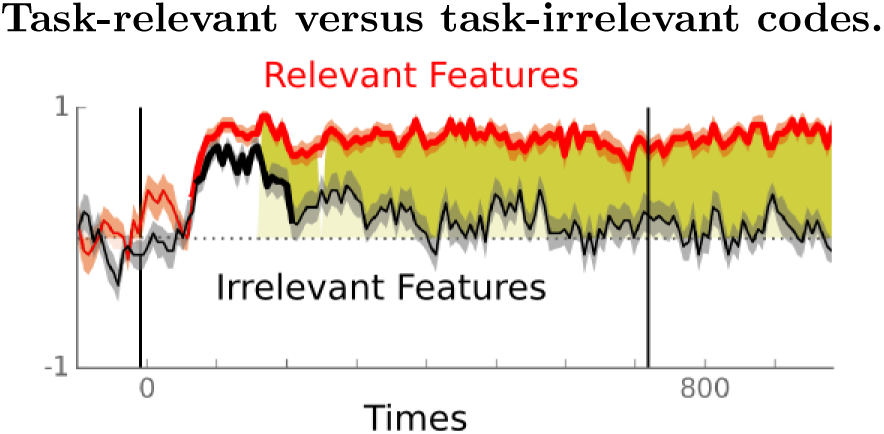
When pulled together, task-relevant decoding scores (target presence, orientation and visibility, in red) can be sustainably decoded from ∼80 ms up to the end of the trial. On the contrary, task-irrelevant features (target spatial frequency, phase and contrast, in black) can only be decoded between ∼80 and 250 ms. Thick lines indicate cluster corrected p < 0.05. Yellow areas indicate when relevant decoding scores are significantly different from irrelevant decoding scores.

**Figure S11:**
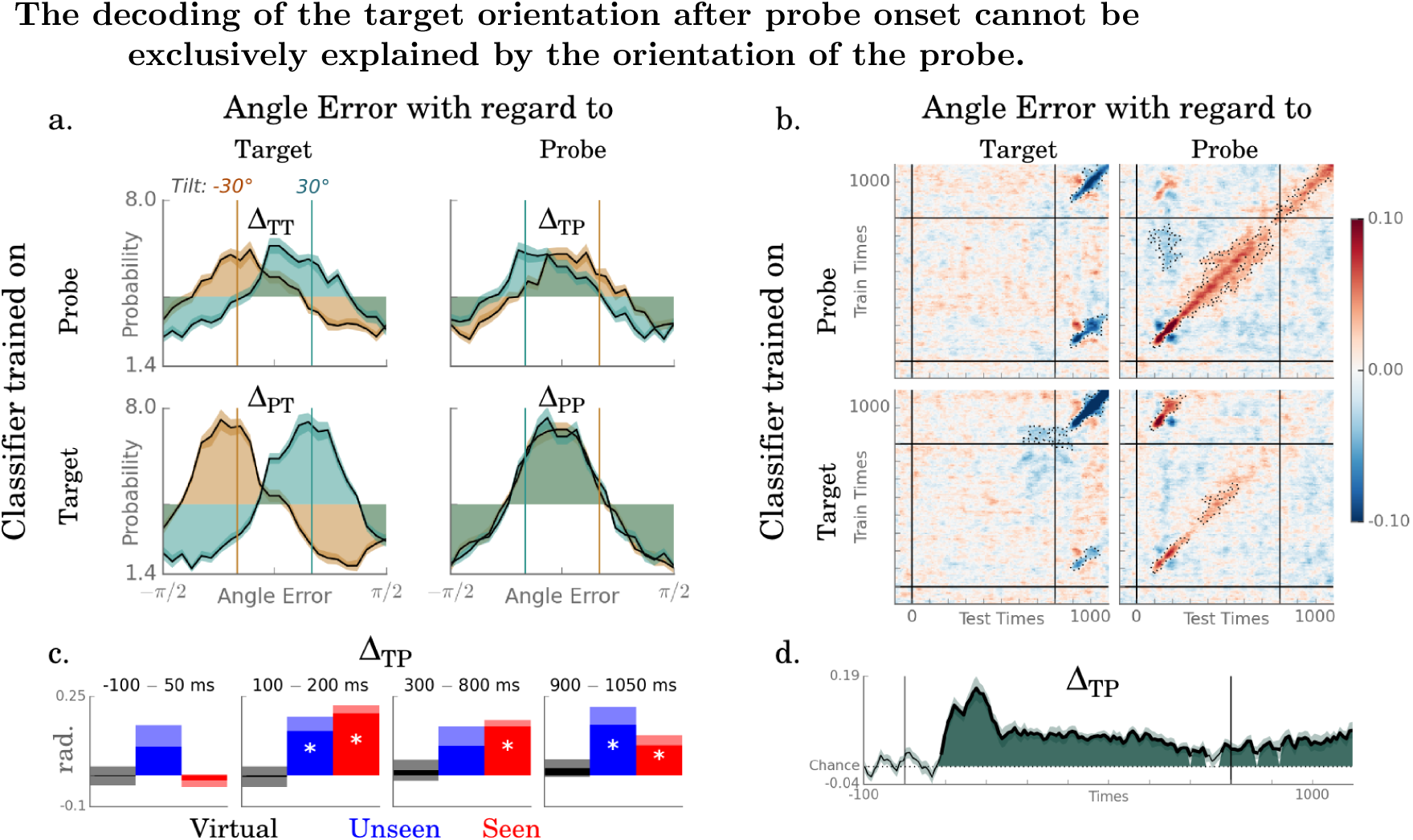
a. Decoding error relative to the target (left) and to the probe (right) for estimators trained on the target (top) or on the probe (bottom) separately for each target-probe tilt condition (brown versus steel blue). The decoding of the target after probe onset (900–1050 ms) is biased toward the probe (top left), but remains correlated with the orientation of the target (top right). These results confirm that both the orientations of the target and of the probe can be decoded after probe onset. b-d. The temporal generalization matrices and the decoding time course of the target-bias effects confirm that we can decode the orientation of the target independently of the probe throughout the entire epoch. c. Target-bias analyses applied separately for seen (red), unseen (blue) and virtual trials (black), confirm that unseen stimulus can be decoded early after probe onset. Error bar indicate SEM across subjects, stars indicate significant (p < 0.05 target-bias effect.)

## 7 Supplementary Materials

### 7.1 Is the decoding of the target orientation after probe onset explained by the orientation of the probe?

Decoding the orientation of the target after the onset of the probe is ambiguous, because the angles of these two stimuli correlated: the probe orientation was either tilted 30^°^ or −30^°^ to the target. The fact that probe estimators did not predict the orientation of the probe (Figure 3), is already indicative that the correlation between the target and probe angles is unlikely to fully account for the target angle decoding scores obtained after probe onset. To formally test this issue, we nevertheless ran a series of control analyses.

#### Method

To test whether the decoding of the target orientation was influenced by the orientations of the target and/or of the probe, we quantified, separately for each prediction of each target angle estimator, the angle errors Δ relative to the target angle (Δ_*TT*_ = *ŷ*_*Target*_ - y_*Target*_) and to the probe angle respectively (Δ_*TP*_ = *ŷ*_*Target*_-*y*_*Probe*_). These angle errors were then compared as a function of the single-trial target-probe tilt). The rational of this analysis stems from a simple principle: if the target angle decoding depends solely on probe angle, then the decoding angle error relative to the probe angle should not vary as a function of target – probe tilt. To test this hypothesis we thus computed the correlation coefficient R^2^ between the angle error obtained in each trial and the target-probe tilt (-1 or 1) with a circular linear correlation. To obtain a signed bias estimate, we computed.

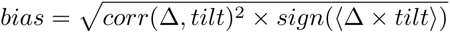
where (Δ x *tilt*) is the mean of the product between the angle error and the direction of the tilt across trials, applied complex space (see method).

This measure of decoding bias thus ranges between [−1,1] and its chance level is 0. Second-order statistical analyses of biases can thus be applied for each decoding analyses of circular data in a similar way to the other analyses of the present study.

Such estimates can thus quantifies the extent to which target angle estimators are biased towards the target relative to the probe (Δ_*TP*_) as well as to the target (Δ_*TT*_). To completely estimate the independence between the decoded angles of the target and of the probe, we also tested how probe angle estimators were biased relative to the target (Δ_*PT*_) and to the probe (Δ_*PP*_).

#### Results

We i) analyzed trials as a function of the probe-target tilt (clockwise or counter-clockwise) and ii) quantified the errors of decoding angle relative to target (Δ_*TT*_) and relative to the probe (Δ_*TP*_) separately. The rational of this analysis is based on the following principle: should the decoding scores be exclusively driven by the probe, the decoding error relative to the probe should be independent of the target-probe tilt.

The results precisely rejected this hypothesis (Figure S11): while the presence of the probe did contribute to the decoding of the target angle, Δ_*TP*_ critically varied as a function of the target-probe tilt, indicating that the decoding error relative to the probe was biased towards the angle of the target. This effect was observed from ∼100 ms after target onset and up to the end of the epoch (100–250 ms: R=0.208±0.006, p<0.001, 300–800 ms: 0.131±0.014, p<0.001, 900–1050 ms: 0.093±0.022, p=0.001). Critically, the effect of Δ_*TP*_ remained significant after probe onset for both seen (R=0.094±0.032, p=0.010) and unseen trials (R=0.161±0.062, p=0.004). To check the validity of this ad-hoc analysis, we also tested it on ‘virtual’ trials. Specifically, we arbitrarily assigned absent trials with a random target probe tilt value, and successfully verified that its decoding error relative to the probe angle (*Δ*_*VP*_) did not vary with the virtual tilt (R=0.023±0.027, p=0.412). We also successfully tested these analyses on the baseline time window of seen, unseen and virtual trials (0–100 ms, all p > 0.05).

Finally, we applied reciprocal analyses to the estimators trained on the probe angle. Interestingly, Δ_*PP*_ did not vary with tilt (R=-0.013±0.012, p=0.2043), and this lack of effect was significantly smaller than Δ_*TP*_ (R=-0.106±0.023, p<0.001). These results indicate that the decoding of the probe angle was not affected by the target and thus shows that the neural representations respectively coding for the target and for the probe are, at least partially, independent and simultaneously encoded in the brain.

## 8 Acknowledgments

This work was funded by the Direction Générale de l’Armement and the Bettencourt-Schueller Foundation to JRK. We are infinitely grateful to Sébastien Marti, Denis Engemann, Alex Gramfort, Valentin Wyart, Virginie Van Wassenhove and the MEG group at Neurospin, as well as MNE-Python and Scikit-Learn communities for their invaluable daily support.

